# A statistical definition for reproducibility and replicability

**DOI:** 10.1101/066803

**Authors:** Prasad Patil, Roger D. Peng, Jeffrey T. Leek

## Abstract

Everyone agrees that reproducibility and replicability are fundamental characteristics of scientific studies. These topics are attracting increasing attention, scrutiny, and debate both in the popular press and the scientific literature. But there are no formal statistical definitions for these concepts, which leads to confusion since the same words are used for different concepts by different people in different fields. We provide formal and informal definitions of scientific studies, reproducibility, and replicability that can be used to clarify discussions around these concepts in the scientific and popular press.

## Main text

Reproducibility and replicability are at the center of heated debates in psychology (*1*) genomics (*2*), climate change (*3*), economics (*4*), and medicine (*5*). The conversation about reproducibility and replicability of science has spilled over into the popular press (*6*, *7*) with major political consequences including new governmental policies and initiatives (*8*) focusing on the reliability of science. These studies have generated responses (*9*), responses to responses (*10*), and ongoing discussion (*11*),(*12*).

Everyone agrees that scientific studies should be reproducible and replicable. The problem is almost no one agrees upon what those terms mean. A major initiative in psychology used the term “reproducibility” to refer to completely re-doing experiments including data collection (*1*). In cancer biology “reproducibility” has been used to refer to the re-calculation of results using a fixed set of data and code (*13*). “Replication” is often used to refer to finding two independent studies that produce a result with similar levels of statistical significance in human genetics(*14*), The same word has been used to refer to re-doing experiments (*15*) and recreating results from fixed data and code(*16*).

These disagreements in terminology seem purely semantic, but they have major scientific and political implications. The recent back-and-forth in the pages of *Science* mentioned above hinged critically on the definition of “replication” with disagreement between the authors about what those terms meant. The press, government officials, and even late night comedy hosts have pointed out “irreproducibility” - defined as the inability to re-create statistical results using fixed data and code - as the fundamental problem with the scientific process. But they use this term to encompass all of the more insidious problems of false discoveries, missed discoveries, scientific errors, and scientific misconduct (*17*). Others have suggested conceptual frameworks to help define these terms (*18*) but have stopped short of a statistical model - which is critical for being precise in discussions of often complicated terms.

To address this major difficulty we need a statistical framework for the scientific process. Typically statistical and machine learning models only formally define the outputs of the scientific process with random variables. The outcome is often designated by **Y** and the covariates of interest by **X**. This framework is limited because it does not allow us to model variation in what the scientists in question intended to study, how they performed the experiment, who performed the experiment, what data they collected, what analysis they intended to perform, who analyzed the data, and what analysis was actually performed. We have created a formal statistical model that includes terms for all of these components of the scientific process (see Supplemental Material). The key steps in this process can also be represented using a simple visual model (Figure 1).

**Figure 1.**
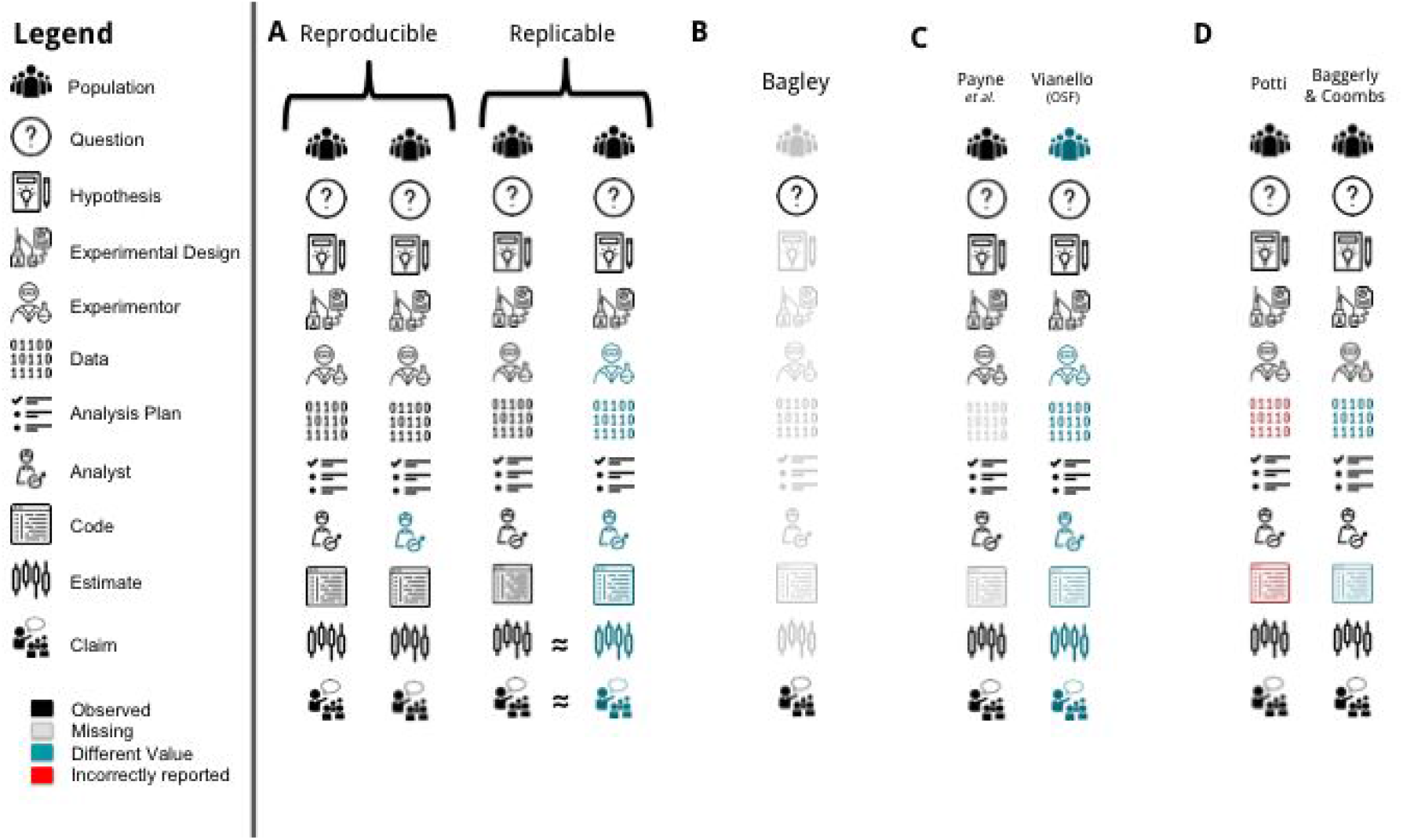
A graphic representation of the statistical model for the scientific process *A*. *Reproducibility is defined as re-performing the same analysis with the same code using a different analyst; replicability is defined as re-performing the experiment and collecting new data. **B**. The paper that only 6 out of 53 pre-clinical studies replicated (*5*) only reported a question and a claim, but not the rest of the scientific components of a study. **C**. The disagreement over the Reproducibility Project Psychology (*9*, *19*) is because a replication was not performed since the population changed. **D**. In the case of the controversy over genomic signatures for chemosensitivity (*2*), reproducibility wasn’t the main issue - the issue was that the original study did not have correct code and data.*

Here we provide informal definitions for key scientific terms and provide detailed statistical definitions in the Using this modeling framework we provide formal definitions for the following terms (see Supplementary Material for formal statistical definitions and additional definitions):

### A scientific study

*consists of document(s) specifying a population, question, hypothesis, experimental design, experimenter, data, analysis plan, analyst, code, parameter estimates, and claims about the parameter estimates.*

### Publication

*Making a public claim on the basis of a scientific study.*

### Reproducible

*Given a population, hypothesis, experimental design, experimenter, data, analysis plan, and code you get the same parameter estimates in a new analysis (**Figure 1A**).*

### Strongly replicable study

*Given a population, hypothesis, experimental design, analysis plan, and code you get consistent estimates when you recollect data and perform the analysis using the original code.*

### Replicable study

*Given a population, hypothesis, experimental design, and analysis plan you get consistent estimates when you recollect data and redo the analysis (**Figure 1A**).*

### Strongly replicable claim

*Given a population, hypothesis, experimental design, analysis plan, and code, you make an equivalent claim based on the results of the study.*

### Replicable claim

*Given a population, hypothesis, experimental design, and analysis plan, you make an equivalent claim based on the results of the study.*

### Conceptually replicable

*A population and a question relate two hypotheses. A scientific study is performed for the first hypothesis and a claim is made. Then a scientific study is performed for a second hypothesis and a claim is made. The claims from the two studies provide consistent answers to the question.*

### False discovery

*The claim at the conclusion of a scientific study is not equal to the claim you would make if you could observe all data from the population given your hypothesis, experimental design, and analysis plan.*

### Garden of forking paths

*Given a population, hypothesis, experimental design, experimenter, data, analysis plan, and analyst, the code changes given the data you observe.*

### P-hacking

*Given a population, hypothesis, experimental design, experimenter, data, analysis plan, and analyst, the code changes to match a desired statement.*

### File drawer effect

*The probability of publication depends on the claim made at the conclusion of a scientific study.*

We can use our statistical framework to resolve arguments and misconceptions around some of the most controversial discussions of reproducibility and replicability. Consider the case of the claim that many of 53 pre-clinical studies were not replicable when performed by scientific teams at a pharmaceutical company (*5*). Under our formal model, the paper describing this replication effort reported a hypothesis - that most studies do not replicate. It also reported a claim - that 47 out of the 53 studies could not be replicated by scientists at the company. However, the population, hypothesis, experimental design, experimenter, data, analysis plan, analysts, code, and estimates are not available (**Figure 1B**). This makes it clear that the published report is missing most of the components of a scientific study.

Later, three replication studies [(*20*), (*21*)/(*22*), (*23*)] were reported from the same pharmaceutical company - though it was unclear if they were part of the originally reported 53. It was pointed out that some of the reported studies included experiments with different populations - violating the definition of a replication. A similar issue was at the heart of a disagreement over several of the studies in the Reproducibility Project: Psychology (*1*). In this project, 100 studies were replicated by independent investigators. In one case, a study originally performed in the United States on US college students was evaluated among a group of Italians. It was pointed out that this change in population violates the definition of replication (*9*) - using our framework it is clear the reason is that the population changed (**Figure 1C**).

Finally consider one of the earliest and most egregious debates over reproducibility - the case of a predictor of chemosensitivity that ultimately fell apart - leading to a lawsuits, an Institute of Medicine conference and report, and ultimately the end of the lead author’s scientific career (*2*). In this case, both the code and the data produced by the original authors were made available; however, they were the wrong code and data. A team from MD Anderson was able to investigate and ultimately produce data and code that reproduced the original results (**Figure 1D**). Ultimately, the study was reproducible, which is surprising given the focus on this study being a violation of reproducibility. The problem with the study was not that the data and code could not be produced, it was that these items, when produced, were wrong (*24*).

Our statistical framework can be used to address other issues that are central to the debate over the validity of science including p-hacking(*25*), the garden of forking paths(*26*), conceptual replication, and other scientific and data analytic problems that arise before the tidy data are processed at the end of the experiment. Using a proper and statistically rigorous framework for the scientific process we can help resolve arguments and provide a solid foundation for journal and public policy around these complicated issues.

